# DNA nanoswitch barcodes for multiplexed biomarker profiling

**DOI:** 10.1101/2020.09.22.309104

**Authors:** Arun Richard Chandrasekaran, Molly MacIsaac, Javier Vilcapoma, Clinton H. Hansen, Darren Yang, Wesley P. Wong, Ken Halvorsen

**Affiliations:** The RNA Institute, University at Albany, State University of New York, New York, NY, USA; Program in Cellular and Molecular Medicine, Boston Children’s Hospital, Boston, MA 02115; Wyss Institute for Biologically Inspired Engineering, Harvard University, Boston, MA 02115; Department of Biological Chemistry and Molecular Pharmacology, Blavatnik Institute, Harvard Medical School, Boston, MA 02115

## Abstract

The detection of molecular biomarkers plays a key role in the clinic, aiding in diagnostics and prognostics, and in the research laboratory, contributing to our basic understanding of diseases. The ability to detect multiple and diverse molecular biomarkers within a single accessible assay would have great utility, providing a more comprehensive picture for clinical evaluation and research, but is a challenge with standard methods. One promising approach is the use of dynamic DNA nanostructures that can respond to molecular biomarkers, which have recently been used in a variety of biosensing strategies. In this work, we report the use of programmable DNA nanoswitches for the multiplexed detection of up to 6 biomarkers within a single pot through the use of a barcoded gel-based readout. We demonstrate the barcoding capability using gene fragments that correspond to 6 different diseases, with each fragment or combination of fragments producing a unique barcode signature. As a defining feature of our method, we show “mixed multiplexing” for simultaneous barcoded detection of different types of biomolecules – DNA, RNA, antibody and protein in a single assay. To demonstrate clinical potential, we show multiplexed detection of a prostate cancer biomarker panel in serum that includes two microRNA sequences and prostate specific antigen (PSA). This strategy holds promise in clinical diagnostics for profiling complex and diverse biomarker panels.

---

Barcodes are ubiquitous in our daily lives, as a way to reduce complex information to a simple pattern. They have also found applications in biosensing, where the study of multiple biological markers can provide useful information for understanding cellular processes as well as disease progression. Some examples of biological barcodes include hydrogel-encapsulated photonic crystals for detection of cardiovascular biomarkers,^[1]^ DNA-antibody conjugates for multiplexed protein analysis,^[2]^ duplex DNA barcodes for cell sorting,^[3]^ DNA-nanoparticle conjugates for nucleic acid^[4]^ and protein detection^[5]^ and DNA-based barcodes for nucleic acid analysis.^[6,7]^ Barcoded architectures can also enable multiplexed detection, where several biomarkers are detected in parallel in a single pot. Such strategies have used conjugated polymers,^[8]^ photonic crystal particles,^[9]^ carbon nanotubes,^[10]^ semiconductor quantum dots,^[11,12]^ DNA-templated silver nanoclusters,^[13]^ gold nanoparticles^[14]^ as well as DNA nanostructures.^[15,16]^ DNA nanostructures in particular are promising for molecular barcodes, as they can be designed to reconfigure in the presence of molecular biomarkers such as proteins, antibodies and nucleic acids. Here, we developed reconfigurable DNA nanoswitches that can be combined to provide barcoded detection and analysis of multiple biomarkers in a single one pot assay (**Figure 1**). Such a system can gather information from multiple types of biomarkers to create a single barcode that can more accurately diagnose a disease, as compared to using only a single biomarker. Studies have already shown that detecting a panel of disease biomarkers is more accurate in diagnosing specific diseases compared to individual biomarkers. For example, biomarker panels that include both the protein biomarker prostate specific antigen (PSA) and microRNAs can outperform diagnosis by PSA testing alone^[17,18]^ and profiling of both SARS-CoV-2 viral antigens and the antibody response in the blood could be useful in tracking and predicting disease progression, such as respiratory failure, in severe COVID-19 cases.^[19]^

**Figure 1.**
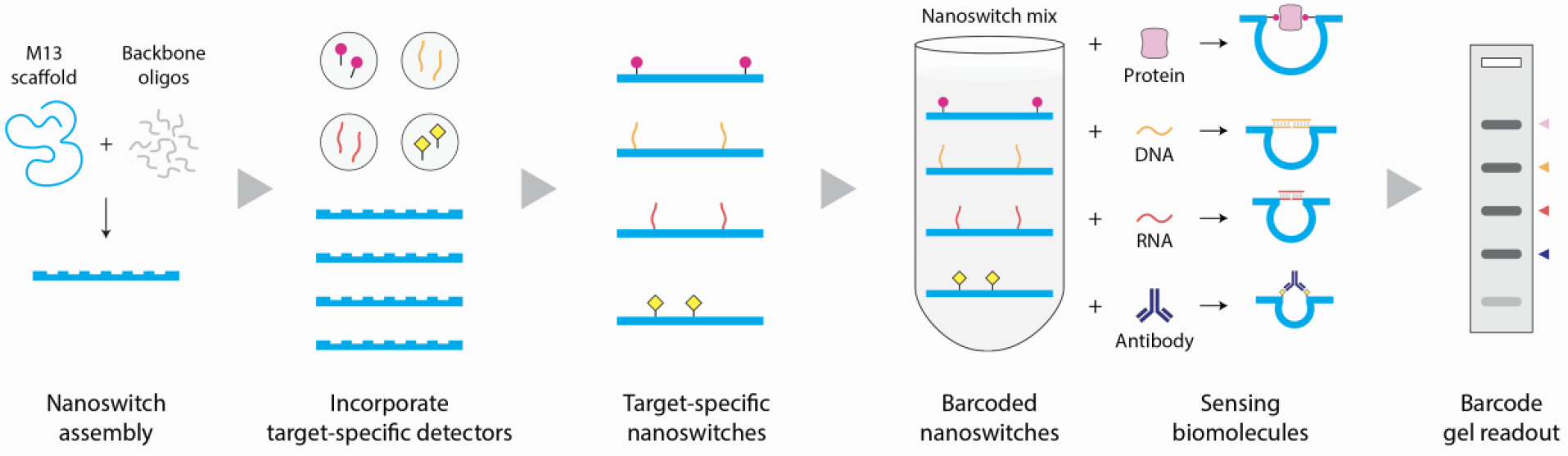
DNA nanoswitch barcodes. Schematic illustration showing construction and use of DNA nanoswitches to yield barcoded results for diagnostic assays.

Programmable DNA nanoswitches are assembled from a long single-stranded scaffold (viral genome M13 routinely used for DNA origami) and short complementary backbone oligonucleotides.^[20]^ Pairs of backbone oligonucleotides can be modified to contain single stranded extensions (detectors) that are complementary to parts of a target nucleic acid (**Figure 2a and S1**). On binding the target sequence, the nanoswitch changes conformation from a linear “off” state to a looped “on” state, providing a distinct signal on an agarose gel (**Figure 2a**, inset). Importantly, this approach requires no complex equipment or enzymatic amplification. The signal comes from the intercalation of thousands of dye molecules from regularly used DNA gel stains (GelRed in this case). We previously used similar DNA nanoswitches for single molecule experiments,^[21]^ detection of microRNAs,^[22]^ viral RNAs,^[23]^ antigens^[24]^ and enzymes,^[25]^ as well as in molecular memory.^[16,26]^ Here, we expand the use of nanoswitches to a multiplexed DNA barcode system that can be used to detect any combination of up to 6 different biomarkers. We further show for the first time that a single barcode can be used to identify different types of biomarkers including proteins, antibodies, DNA and RNA, with clinical potential shown by detecting a prostate cancer biomarker panel in serum that includes two microRNA sequences and prostate specific antigen (PSA).

**Figure 2.**
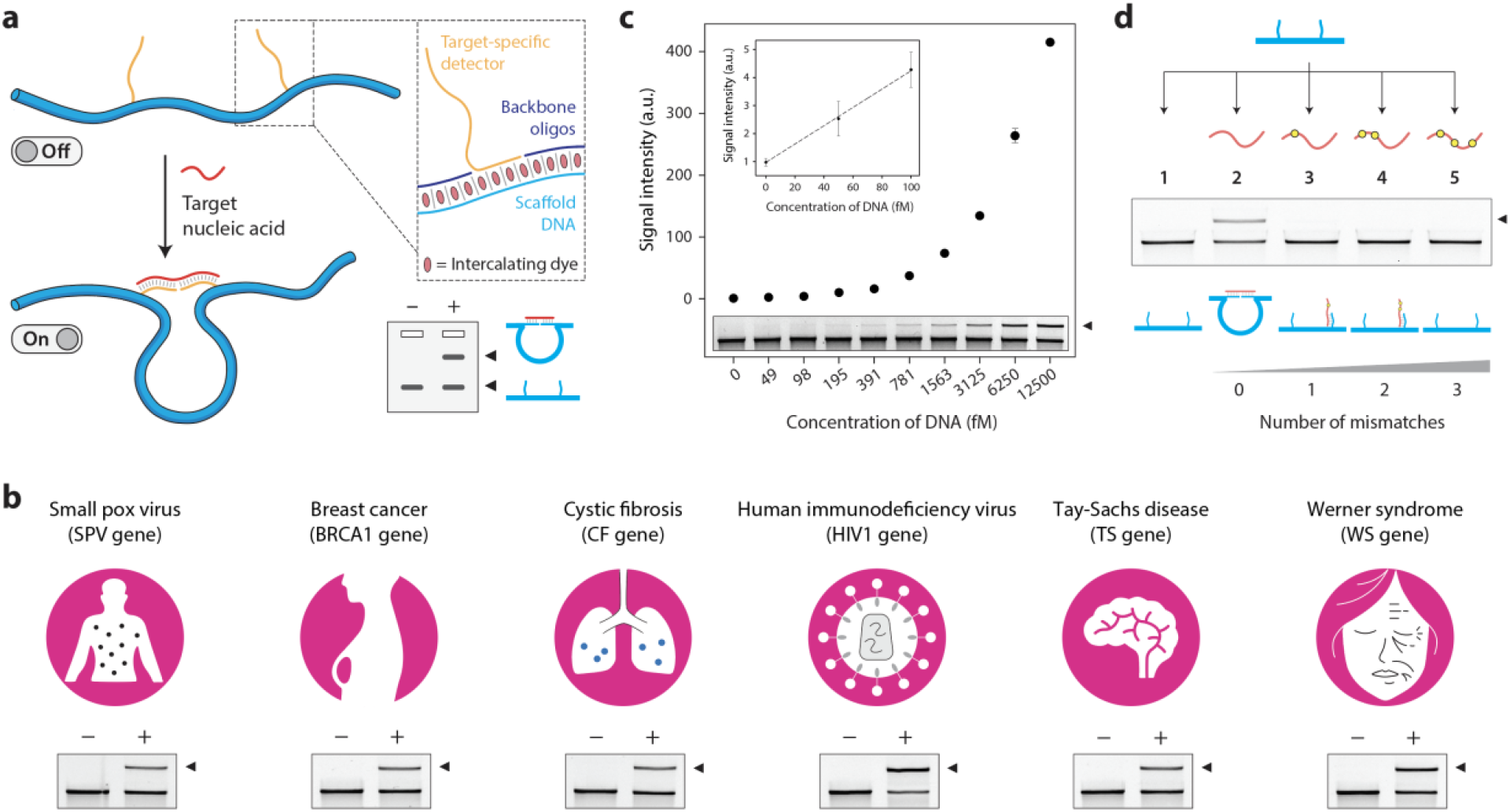
DNA nanoswitch overview and characterization. (a) Reconfiguration of the DNA nanoswitch in the presence of a nucleic acid target turning from a linear “off” state to a looped “on” state. The two states of the nanoswitch can be readout on an agarose gel (inset). (b) Demonstration of detecting different gene fragments using the DNA nanoswitch assay. (c) Sensitivity analysis of DNA nanoswitch assay for the cystic fibrosis (CF) gene fragment. (d) Specificity of DNA nanoswitches against CF gene fragments containing one to three mismatches.

To demonstrate the concept of DNA nanoswitch barcodes, we chose 6 different gene fragments corresponding to the smallpox virus gene (SP), cystic fibrosis gene (CF), Tay-Sachs disease gene (TS), breast cancer gene (BRCA1), human immunodeficiency virus gene (HIV1) and Werner syndrome gene (WS). We designed nanoswitches specific to these gene fragments and showed successful detection of a single-stranded DNA oligonucleotide corresponding to the gene in each case (**Figure 2b and S2**). For further characterization of DNA nanoswitch detection, we chose the cystic fibrosis gene fragment. In diagnostics, sensitivity is a key parameter to detect early onset of biological or disease processes. We performed sensitivity experiments with decreasing concentrations of the DNA and found that the signal could be seen by eye at concentrations as low as 50 fM (**Figure 2c and S3**). Calculating the limit of detection (LOD), defined as the concentration of biomarker that yields a signal that exceeds the mean background by 3 SDs of the background, we obtained a value of ∼12 fM, similar to what we previously reported for protein detection.^[24]^ We then tested specificity of the assay by challenging the CF nanoswitch with CF gene targets that contained 1-3 mutations. Results showed that for a single nucleotide mismatch, there was a 40% reduction in signal compared to the fully complementary target (**Figure S4**). In this assay, the nanoswitch detectors were designed to recognize the complete length of the target sequence (24-nucleotide target hybridized to two 12-nucleotide detectors). To optimize the specificity of the assay, we redesigned the nanoswitch by decreasing the length of the detector complementary to the side of the target that contains the mismatch. Using this design, we showed that the assay is highly specific, able to discriminate even a single nucleotide mismatch in the target sequence (**Figure 2d and Figure S4**). Next, we performed a time series experiment, showing that the assay can be performed in under an hour in most cases (**Figure S5**).

To demonstrate barcoded detection, we used the programmability of the nanoswitch to place the detector strands along specific positions on the scaffold (**Figure 3a**). Nanoswitches with detectors spaced far apart will yield a longer loop on binding the target while a shorter loop will be formed when the detectors are closer together. This allows the creation of specific nanoswitches that yield different bands on the gel according to the resulting loop size (**Figure 3b**). For convenient construction, we designed 12 variable regions (V1-V12 in Figure 3a) for placement of detectors. We designed six nanoswitches with different separations of the two detectors along the scaffold. Each of these nanoswitches contained detectors specific to one of the six gene targets described in Figure 2b. We then prepared a mixture of these nanoswitches that can detect all six targets simultaneously: each target will trigger the specific nanoswitch and form the corresponding loop, providing a unique signature on a gel, i.e. a barcode. The separation of the detectors was chosen in a way that the six looped nanoswitches can be easily resolved on a single gel lane (**Figure 3b, inset**). Before we tested simultaneous detection of the gene fragments, we confirmed that each individual nanoswitch does not have any cross-reactivity with non-specific targets (**Figure 3c**). Next, we used the nanoswitch mixture and demonstrated detection of all possible combinations of the six gene targets (a total of 64), each providing a unique barcode (**Figure 3d and S6**). This strategy provides a simple multiplexed assay to detect several nucleic acids in a single assay, with each recognition event translated into a unique readout. Migration of these nanoswitches in the gel is dependent primarily on the size and location of the loop, rather than on the molecular weight of the target strand (since the target strand is only 20-30 nucleotides compared to the ∼7 kbp nanoswitch). Thus, the barcodes generated by the different targets is constant regardless of the target of interest, opening up the possibility of detecting different types of biomarkers simultaneously and not just nucleic acids.

**Figure 3.**
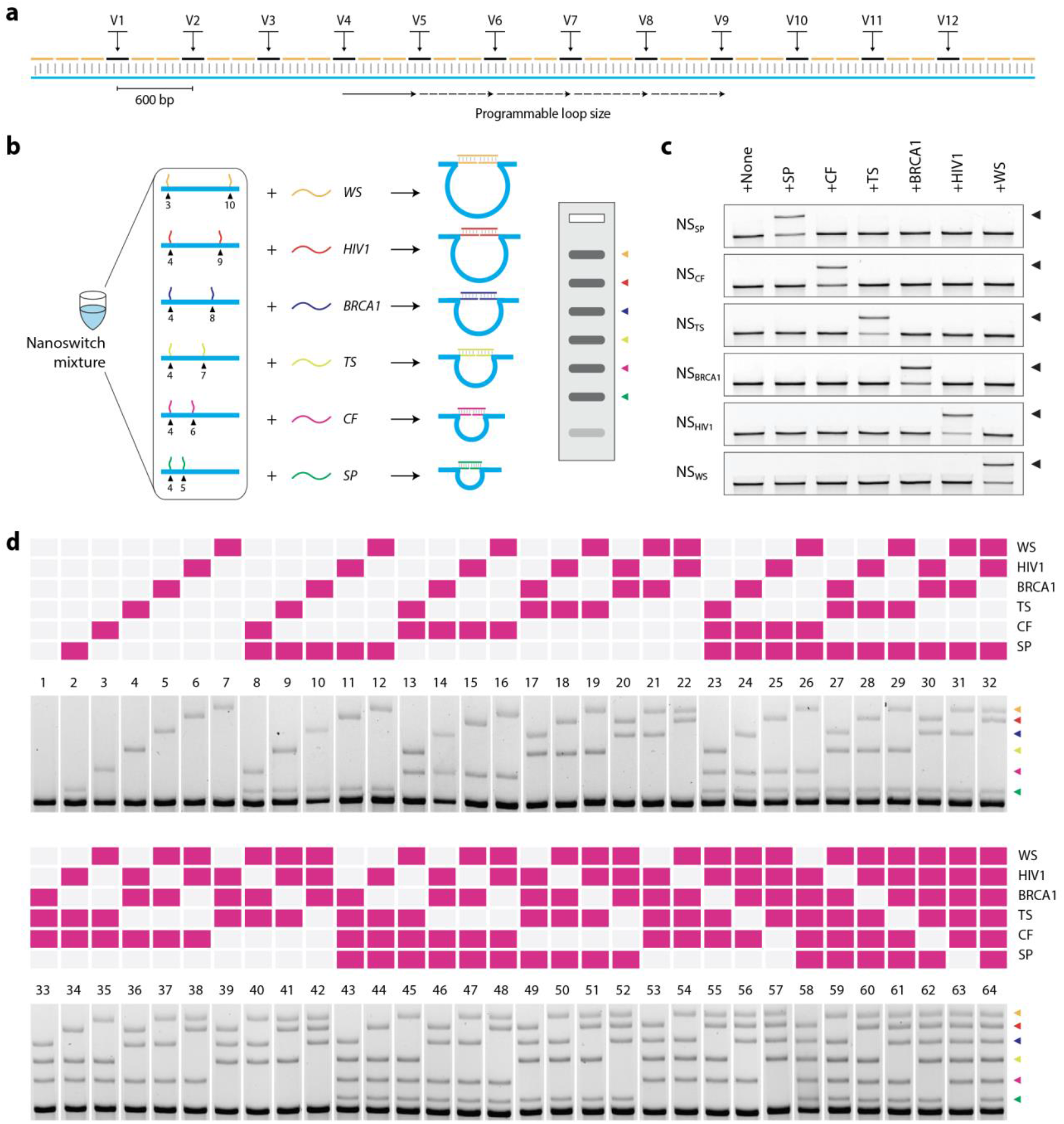
DNA nanoswitch barcodes for multiplexed detection of nucleic acids. (a) Design of the DNA nanoswitch with variable regions (V1-V12) that allow programmable placement of detectors, resulting in nanoswitches with different loop sizes. (b) A mixture of nanoswitches with different loop sizes yields a multiplexed gel readout of up to 6 gene fragments (inset). Each target nucleic acid triggers the formation of a specific loop, providing a unique signal (targets are colored for clarity). (c) Demonstration of lack of cross-reactivity in nanoswitches designed for 6 different gene fragments. (d) Full set of DNA nanoswitch barcodes showing detection of all possible combinations of the 6 gene fragments.

We then extended the nanoswitch barcode design for different types of targets including proteins, antibodies, DNA and RNA. For proof-of-concept protein detection (streptavidin), we designed a nanoswitch where the detector strands were modified to contain a biotin group instead of single - stranded extensions. Similarly, for detecting an antibody (anti-digoxygenin), we incorporated digoxygenin-coupled detectors in the nanoswitch (**Figure 4a-b**). We designed two more nanoswitches that can detect specific DNA and RNA sequences. We also designed the four nanoswitches to yield different loop sizes so that a nanoswitch mixture can provide barcoded recognition for all four targets in a single assay (**Figure 4c**). Again, we first confirmed that there was no cross-reactivity between these nanoswitches when tested against non-specific targets (**Figure 4d**). We then showed all possible detection events (16 in total for 4 targets) using this barcode (**Figure 4e and S7**), demonstrating that our approach can simultaneously detect nucleic acids, proteins and antibodies in a single one-pot assay. Further, to show that the individual biomarkers can be quantified, we performed a “multiplexed non-interference” analysis by changing the concentration of one biomarker while keeping the others constant. We show that the concentration series for each of the four biomarkers (anti-digoxygenin antibody, DNA, RNA and streptavidin) can be monitored in the presence of other targets (**Figure 4f and S8**). These results are consistent with those where the targets are present alone in a reaction when detected using the same nanoswitch mix (**Figure S9**).

**Figure 4.**
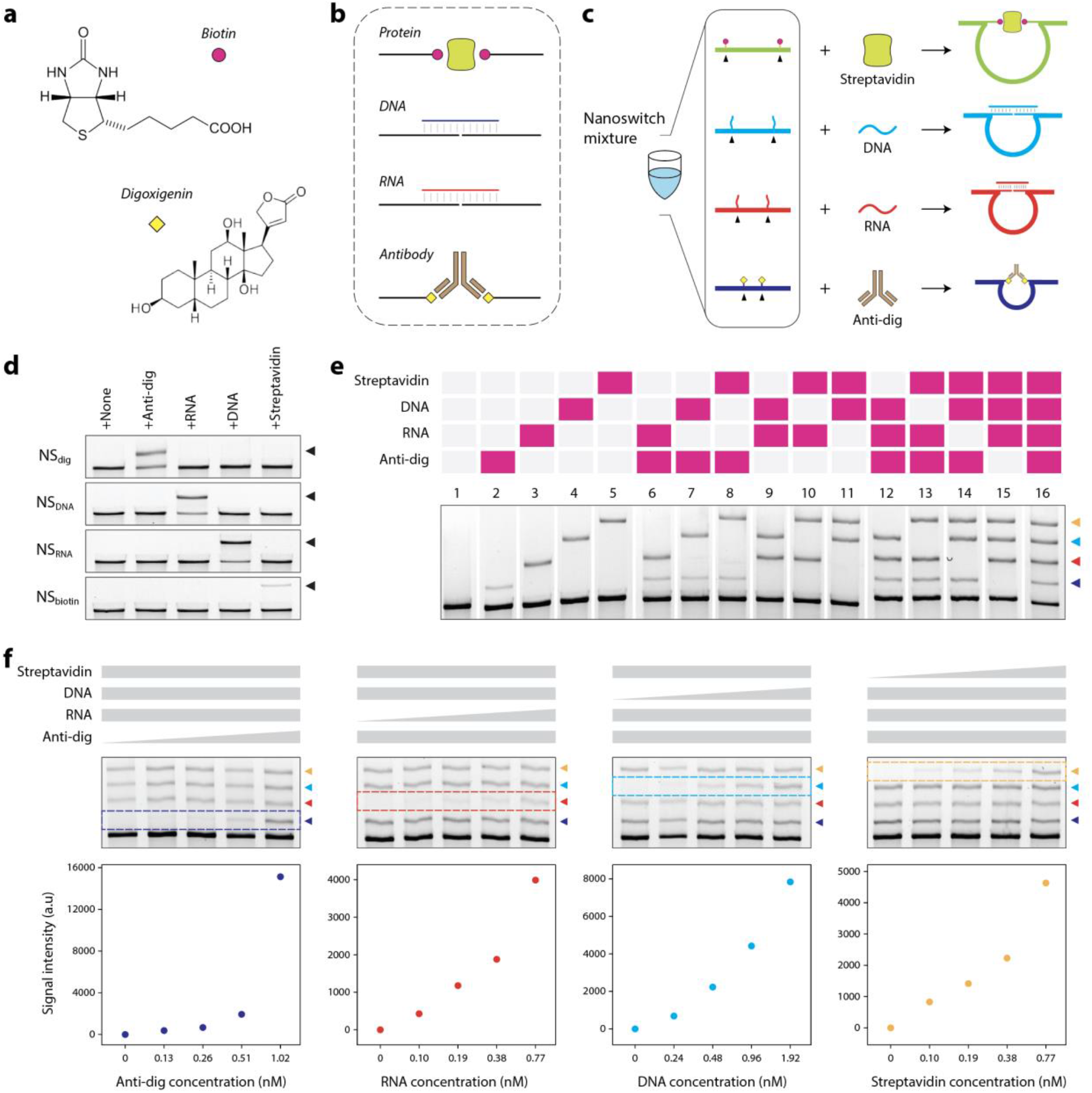
Detecting multiple types of biomarkers using DNA nanoswitch barcodes. (a) Using small molecules such as biotin and digoxygenin to target macromolecules. (b) Strategy to detect nucleic acids (DNA and RNA), a protein (streptavidin) and an antibody (anti-digoxygenin). (c) A nanoswitch mixture to demonstrate simultaneous detection of four different biomarkers. (d) Demonstration of lack of crosstalk in nanoswitches designed for four different target biomolecules. (e) Full set of DNA nanoswitch barcodes showing detection of all possible combinations of anti-digoxygenin (anti-dig), an RNA sequence, a DNA sequence and streptavidin. (f) Individual targets can be analyzed even in the presence of other target biomolecules, showing a “multiplexed non-interference” in the barcoded assay.

Toward clinical relevance of the diagnostic barcodes, we then chose to detect a biomarker panel for prostate cancer (**Figure 5**). Detecting a panel of disease biomarkers rather than an individual biomarker can provide additional information for more accurate diagnoses.^[18]^ For example, data suggests that biomarker panels that include PSA, miR-141 and miR-30c can outperform diagnosis by PSA testing alone.^[17]^ In the multiplexed barcode demonstration above, we showed the detection of proteins and antibodies using small molecule ligands. For detecting PSA, we modified the nanoswitch detectors to contain PSA-specific antibodies. This strategy of using nanoswitches to detect antigens (which we called NLISA, nanoswitch-linked immunosorbent assay^[24]^) is similar to sandwich ELISA where a pair of antibodies are used to detect proteins of interest. In addition to PSA, we included DNA analogs of the microRNA sequences miR-30c and miR-141 based on literature demonstrating the utility of microRNA biomarkers in blood for detection of prostate cancer.^[28,29]^ To detect this panel, we designed three nanoswitches with different loop sizes and created a multiplexed nanoswitch mixture and tested detection of all biomarkers in buffer (**Figure 5c**). We then spiked the biomarkers in 20% serum and showed detection of individual biomarkers (**Figure 5c**, lanes 2-4) or simultaneous detection of all three biomarkers in a single gel lane (lane 5).

**Figure 5.**
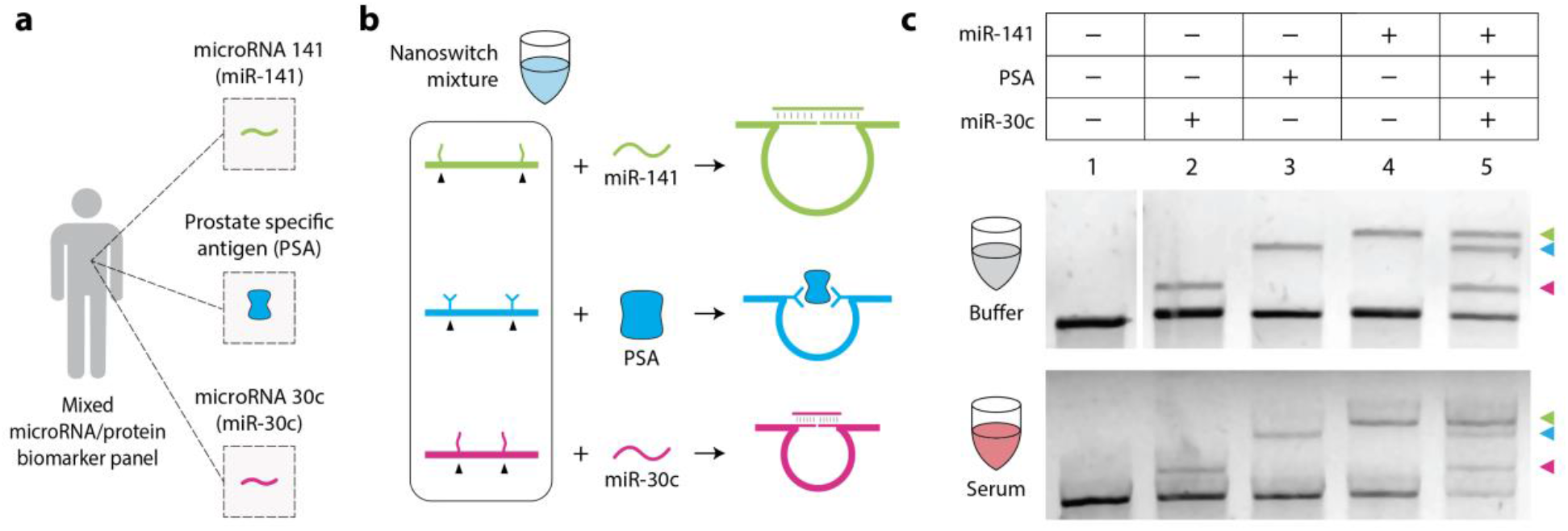
DNA nanoswitch barcodes for prostate cancer biomarker panel. (a) An illustration of the three chosen prostate cancer biomarkers (miR-141, miR-30c and PSA). (b) Barcode detection of individual biomarkers (DNA analogs of microRNAs 141 and 30c, and PSA) and simultaneous detection of all three biomarkers in buffer and 20% fetal bovine serum (FBS).

In this study, we showed that DNA nanoswitch barcodes can be used to detect up to six biomarkers in a single assay, and even a mixture of protein, antibody and nucleic acids. Furthermore, we demonstrated multiplex detection of protein and nucleic acid biomarkers from serum. The nanoswitches can be dried and stored for later use, and are stable after drying and retain their functionality to provide detection barcodes (**Figure S10**). Conveniently, our strategy uses gel electrophoresis for readout, which can be conducted outside of a laboratory setting, using, for example, an electronic buffer-less gel system (ThermoFisher). The barcode system described here can also be combined with nanopore^[28]^ or microfluidic chip^[29]^ based readouts or in sensor arrays for macroscopic graphical readouts.^[30]^ The barcoded assay described here can discriminate even a single mismatch in the target sequence, which is an important parameter for use in detecting single nucleotide polymorphisms (SNPs).^[31]^ Further, unique to our barcoded assay, a variety of biomarkers can be detected in a single pot with a common consolidated workflow, making this strategy useful for detecting biomarker panels, particularly when splitting the sample up would be unfeasible due to challenges of efficiency and sensitivity.

## SUPPORTING INFORMATION

Supporting information can be found online.

## ACKNOWLEDGEMENT

Research reported in this publication was supported by the NIH through NIGMS under award R35GM124720 to K.H. and NCI under award R21 CA212827 to W.P.W. and K.H. Additional funding was provided by the Boston Children’s Hospital Technology Development Fund (W.P.W.)

## AUTHOR CONTRIBUTIONS

A.R.C., K.H. and W.P.W conceived the project. A.R.C designed experiments. A.R.C., J.V., M.M., C.H.H., and D.Y. performed experiments. A.R.C. analyzed and visualized data and wrote the first draft of the manuscript. A.R.C., C.H.H., W.P.W. and K.H. supervised the project. All authors contributed to and edited the final manuscript.

