## Supplementary Information for "DNA nanoswitch barcodes for multiplexed biomarker profiling"

### MATERIALS AND METHODS

#### Materials

Oligonucleotides (including RNA, biotin-modified DNA and digoxygenin-modified DNA) were purchased from Integrated DNA Technologies (IDT) with standard desalting. M13 circular DNA and BtsCI enzyme were purchased from New England Biolabs (NEB). GelRed nucleic acid stain was purchased from Biotium, Fremont, CA, USA. Molecular biology grade agarose was purchased from Fisher BioReagents.

#### Linearization of M13 DNA

5 μl of 100 nM circular single-stranded M13 DNA, 2.5 μl of 10× Cut Smart buffer, 0.5 μl of 100 μM BtsCI restriction-site complementary-oligonucleotide and 16 μl of deionized water were mixed and annealed from 95 °C to 50 °C in a T100™ Thermal Cycler (Bio-Rad, Hercules, CA, USA). 1 μl of the BtsCI enzyme (20,000 units/ml) was added to the mixture and incubated at 50 °C for 15 min. The mixture was brought up to 95 °C for 1 min to heat deactivate the enzyme followed by cooling down to 4 °C.

#### Antibody coupling

Antibodies were coupled to oligonucleotides as previously described.[24] In brief, sandwiching antibodies against PSA (Medix Biochemica Anti-h PSA 8301 and Anti-h PSA 8311) were buffer exchanged in 1× PBS using two Zeba columns (Thermo Fisher Scientific) to remove storage sodium azide. 60 μL of washed PSA antibody at ~4.0 μM were mixed with an equimolar amount of DBCO-PEG4-NHS ester linker (Sigma) in 40 mM, pH 8.6 HEPES buffer and incubated at room temperature for 30 min. Then, 5-fold excess of azide-modified oligonucleotides (Integrated DNA technologies) were added and incubated for 90 min at room temperature. Antibody–oligo conjugates were then purified from excess oligonucleotides and linkers using the Thunder-Link Conjugate Clean Up Reagent (Innova Biosciences) and resuspended in 40 μL of 1× PBS. Antibody coupling and purification were confirmed using 4-20% TBE polyacrylamide gels (Bio-Rad) stained with Krypton Fluorescent Protein Stain (Thermo Fisher Scientific).

**Construction of nanoswitches**

For genotyping, biotin and digoxygenin nanoswitches, linearized single-stranded M13 DNA (20 nM) was mixed with 10-fold excess of the backbone oligonucleotides and detector strands and annealed from 90 °C to 20 °C at 1 °C $min^{-1}$ in a thermal cycler. Following construction, the nanoswitches were purified using HPLC to remove excess oligonucleotides or used unpurified after dilution in 1× PBS to a concentration of 400 pM. For PSA nanoswitches, linearized single-stranded M13 DNA (20 nM) was mixed with 10-fold excess of the backbone oligonucleotides and annealed from 90 °C to 20 °C at 1 °C $min^{-1}$ in a thermal cycler. 80-fold excess of the conjugated antibody-oligonucleotide detectors were added when the hybridization protocol reached 37 °C. Following construction, the nanoswitches were purified with BluePippin using 0.75% agarose, 1-50kb gel cassette (Sage Science) to remove excess oligonucleotides and antibody-oligo conjugates. The PSA nanoswitches were stored undiluted in protein lobind tubes (Eppendorf) at 4 °C.

**Nanoswitch characterization**

A typical reaction contained 160 pM nanoswitch with 2.5 nM DNA targets in a 10 μl reaction. Samples were incubated at room temperature for 1 h. For sensitivity experiment, the target DNA was spiked into a 500 nM solution of off-target "blocking" oligos to minimize loss to the tubes. Microcentrifuge tubes and pipette tips were additionally pre-incubated in blocking oligo solution to minimize loss of the target DNA. The nanoswitch/DNA target solution was incubated overnight at room temperature in a solution containing 1× PBS and 10 mM $MgCl_2$. For specificity experiments, the final concentration of target strands were 2.5 nM and incubation time was 1 h at room temperature in a solution containing 1× PBS.

**Nanoswitch barcode operation for gene identification and biomarker detection**

Nanoswitch mixture was prepared by mixing equimolar amounts of individual nanoswitches (6 nanoswitches for multiplexed gene analysis and 4 nanoswitches for mixed multiplexing). For barcoded detection, a typical reaction contained final concentrations of ~160 pM nanoswitch mixture, 1× PBS and 2.5 nM DNA gene fragments. For barcodes of mixed biomarkers, typical concentrations were ~1 nM anti-digoxygenin, ~0.8 nM RNA, ~2 nM DNA and ~0.8 nM streptavidin. For concentration series, different concentrations of the biomarkers were added as indicated in Figure 4 and Figure S9.

**Biomarker panel detection in serum**

Nanoswitch mixture was prepared by mixing miR-30c, PSA, and miR-141 nanoswitches to a final molar ratio of 1:2:3 respectively. EDTA was added to 20% FBS to a final concentration of 100 mM. For serum detection, a typical 11 μL reaction contained 6 μL of 20% FBS with 100mM EDTA, biomarkers at a final concentration of 2.5 nM, 1× PBS, and 2 μL of nanoswitch mixture (final concentrations of 30 pM, 60 pM, and 90 pM). The biomarkers used were PSA antigen (Biospacific, cat#: J63011, high purity) and ssDNA sequences of miR-141 and miR-30c (Integrated DNA technologies). Samples were incubated at room temperature for 90 minutes.

**Gel electrophoresis**

Nanoswitches were run in 0.8% agarose gels, cast from molecular biology grade agarose dissolved in 0.5× Tris-borate EDTA (TBE). Characterization gels of individual nanoswitches were typically run at 75 V (constant voltage) at room temperature and barcode gels (multiplexing) were run at 55 V (constant voltage) at 4 °C. Samples were pre-stained by mixing 1× GelRed stain and a Ficoll based loading dye (15% Ficoll, 0.1% bromophenol blue) with the samples before loading. 10 μl of the samples were loaded and run on an unstained agarose gel. Gels were imaged with a Bio-Rad Gel Doc XR+ gel imager and analyzed using ImageJ. For the prostate cancer biomarker panel, nanoswitches were run in 0.8% agarose gels at 70 V (constant voltage) at room temperature. After incubation, samples were diluted by adding 1× volume of 0.5× TBE. Samples were then pre-stained with 1× GelRed stain and a Ficoll based loading dye (Promega) and 0.25 μL of 1% Coomassie Brilliant Blue G-250 were added before loading. Gels were imaged with an Invitrogen iBright FL1000 Imaging System and analyzed using ImageJ. Median filter in ImageJ was used to remove noise in gel images in Figures 5b and S10.

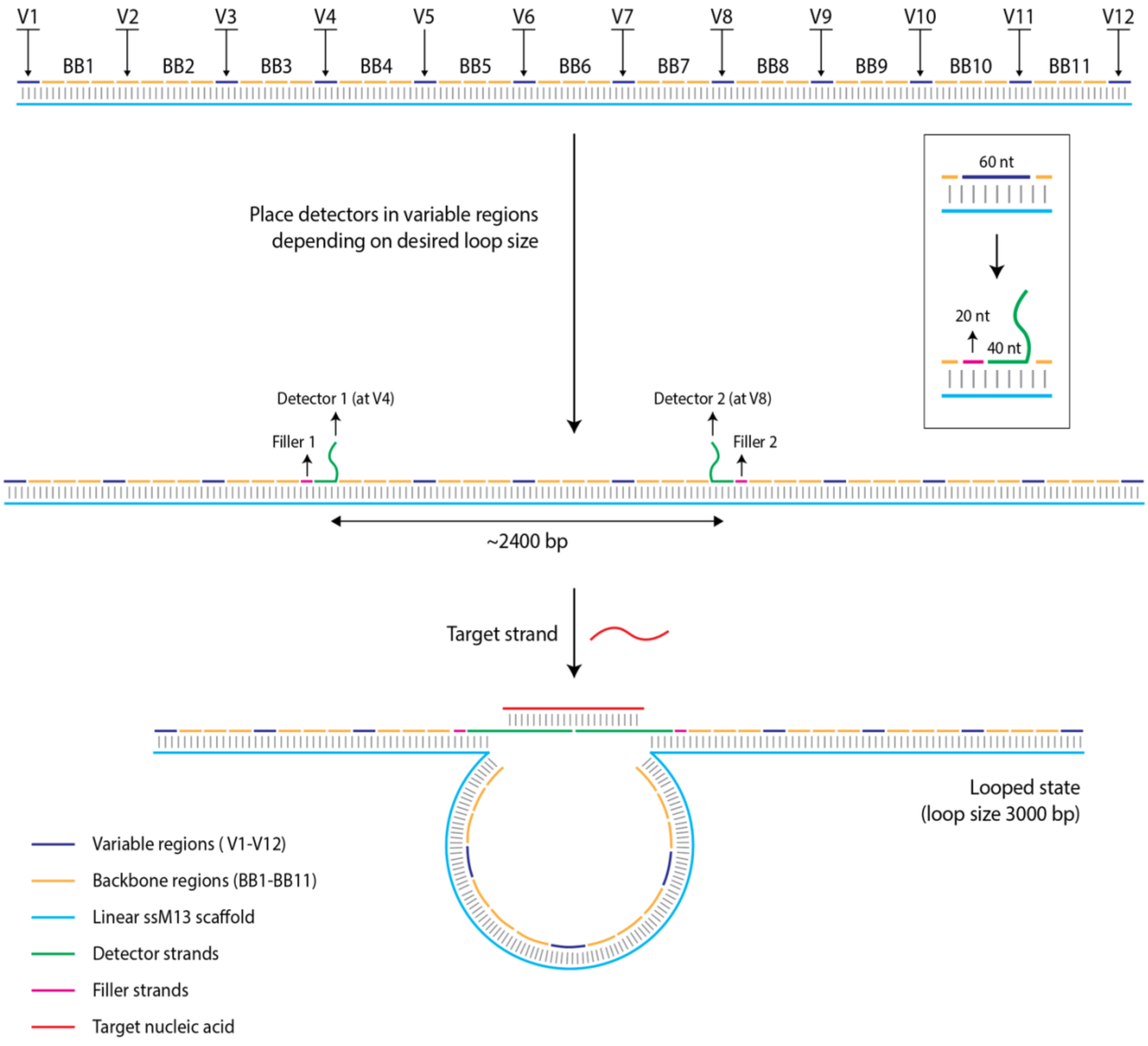


**Figure S1.** The nanoswitch is a duplex formed from linear M13 and short complementary backbone oligonucleotides. Twelve regions (60 nt each) are designated as “variable” regions. Two detectors containing single-stranded overhangs that complement the target can be inserted in place of two of the variable regions. The distance between the two detectors dictates the loop size and migration of the looped state on a gel.

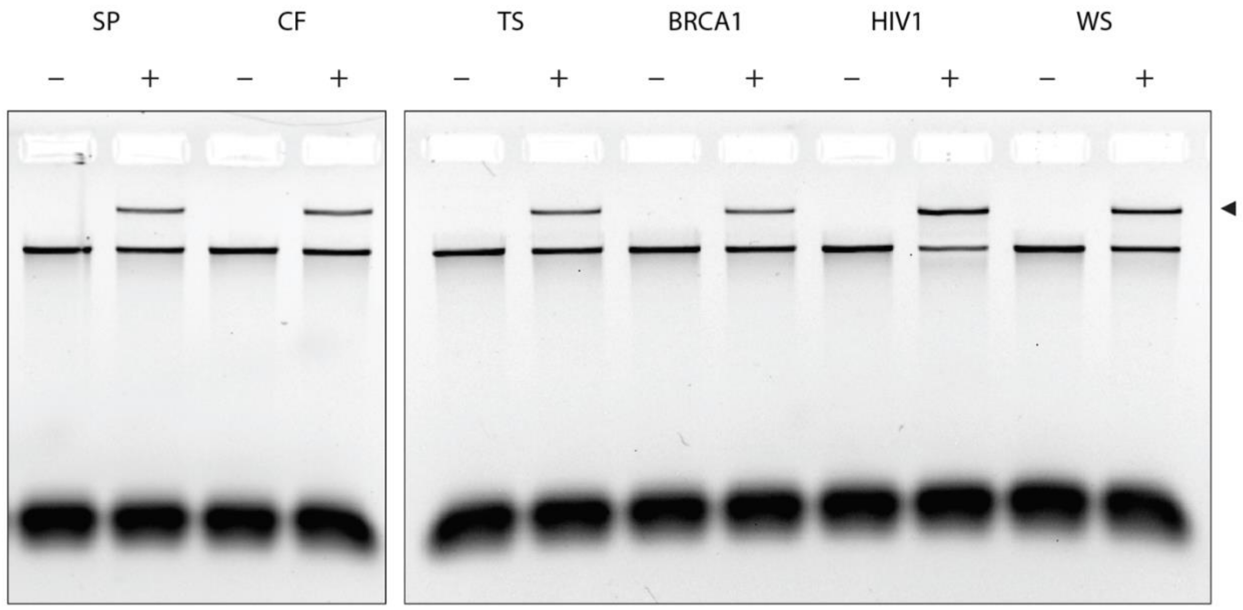


**Figure S2.** Detection of different gene fragments using DNA nanoswitches. Gel shown here is the full image of the gel shown in Figure 2b.

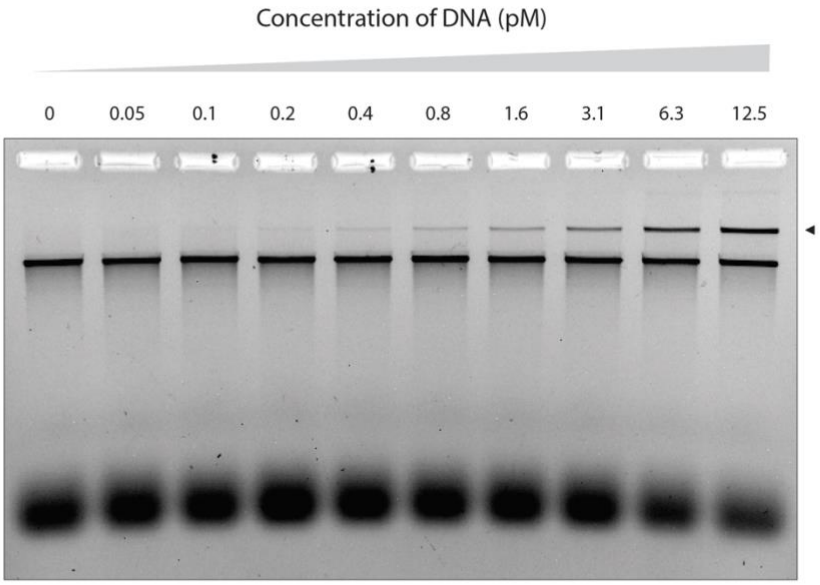


**Figure S3.** Sensitivity of DNA nanoswitch assay for cystic fibrosis gene fragment. Full image of gel shown in Figure 2c.

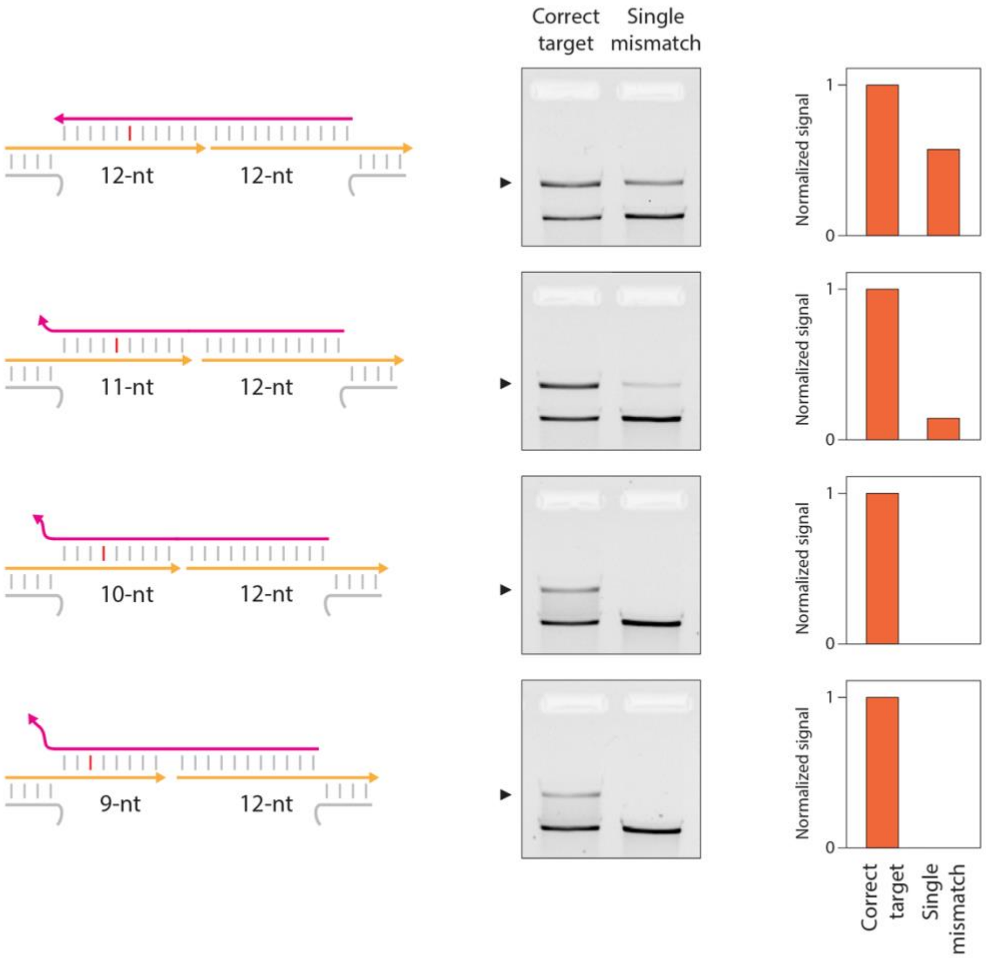


**Figure S4.** Specificity of DNA nanoswitch assay demonstrated with cystic fibrosis gene fragment. With detector length that complements the whole target sequence, a single mismatched target yielded 40% signal compared to a correctly matched target. On reducing the length of the detector, the specificity to discriminate a single mismatch increased, with no signal for the mismatched target using a 10-nt and 9-nt detector length. The 9-nt detector length was used for the specificity image shown in Figure 2d.

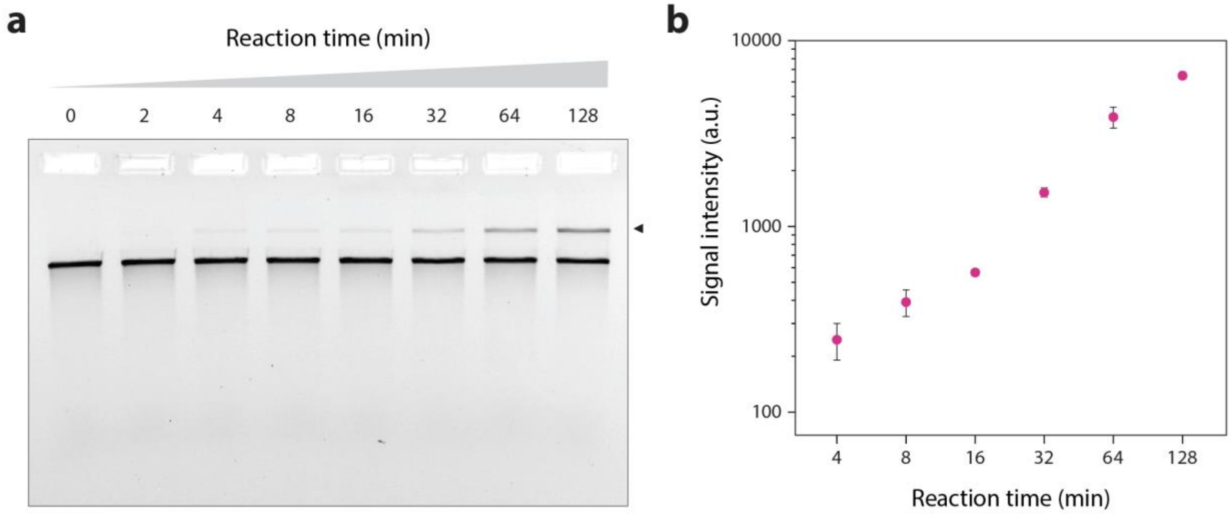


**Figure S5.** Time series of detection of a 1 nM cystic fibrosis gene fragment.

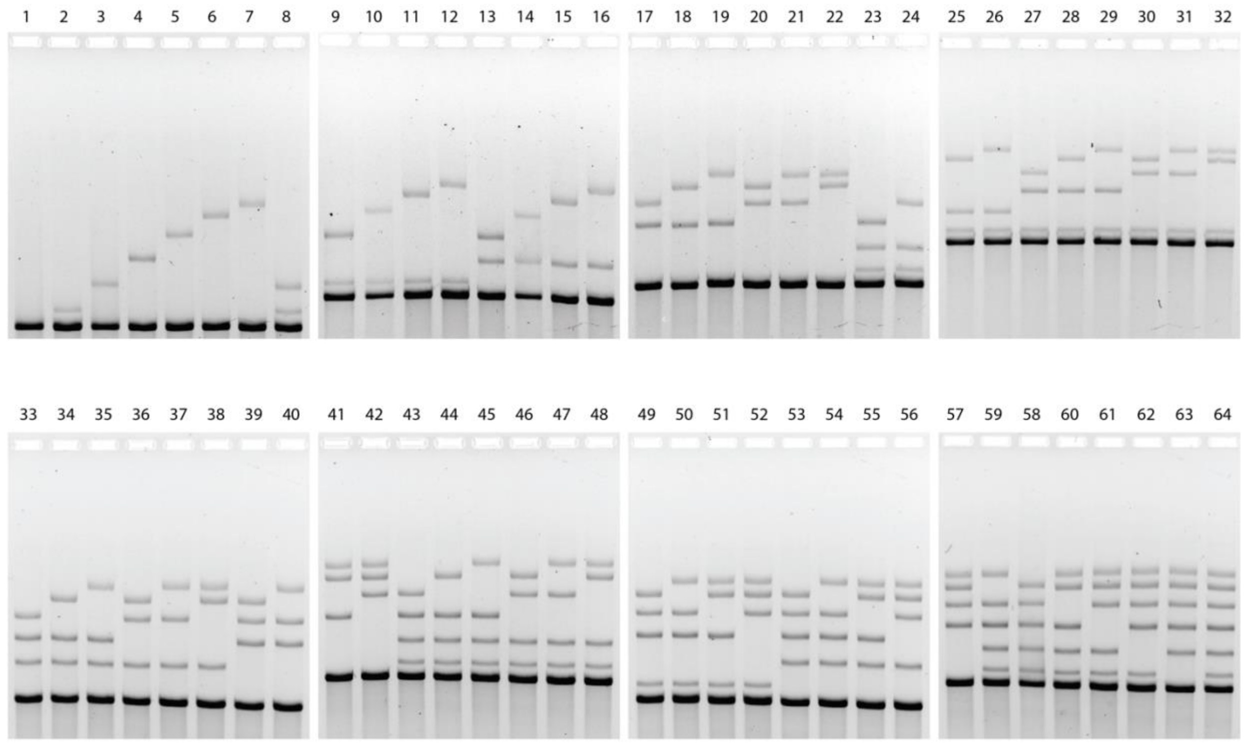


**Figure S6.** Full set of DNA nanoswitch gene barcodes. Gel images show all combinations of six different gene fragments. Full gel images of results shown in Figure 3d.

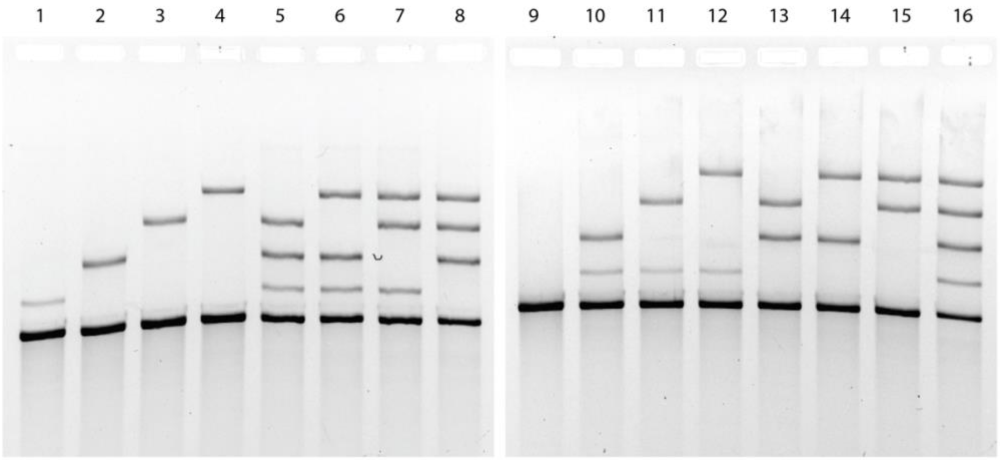


**Figure S7.** Full set of DNA nanoswitch barcodes for multiplexed detection of different types of biomarkers (anti-digoxygenin antibody, RNA, DNA and streptavidin). Full gel images of results shown in Figure 4e.

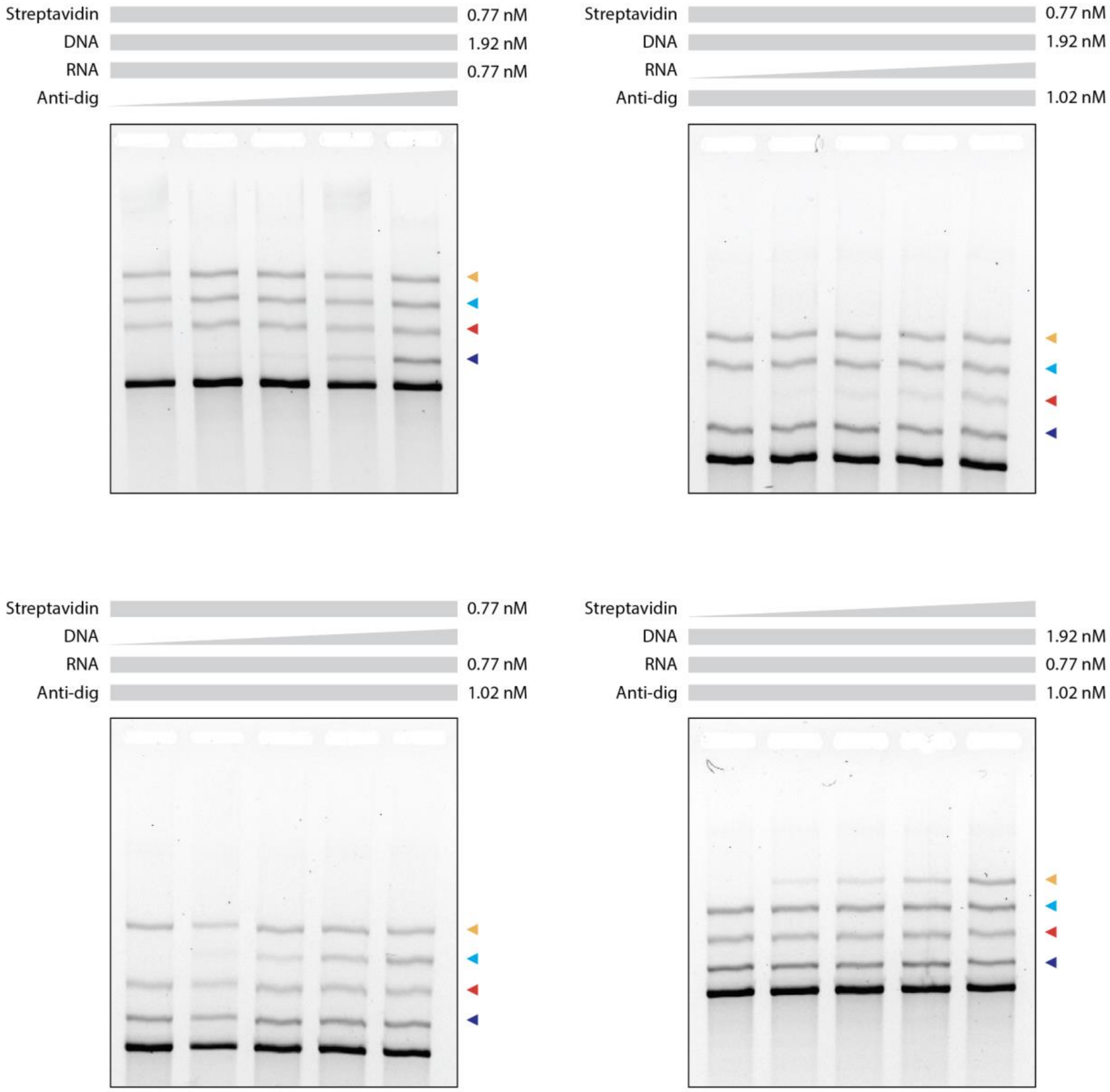


**Figure S8**. Multiplexed concentration series of different biomarkers. Full gel images of results shown in Figure 4f. Constant concentrations of target molecules are indicated above the gel images. The varying concentrations of a single target is shown in Figure 4f and Figure S9.

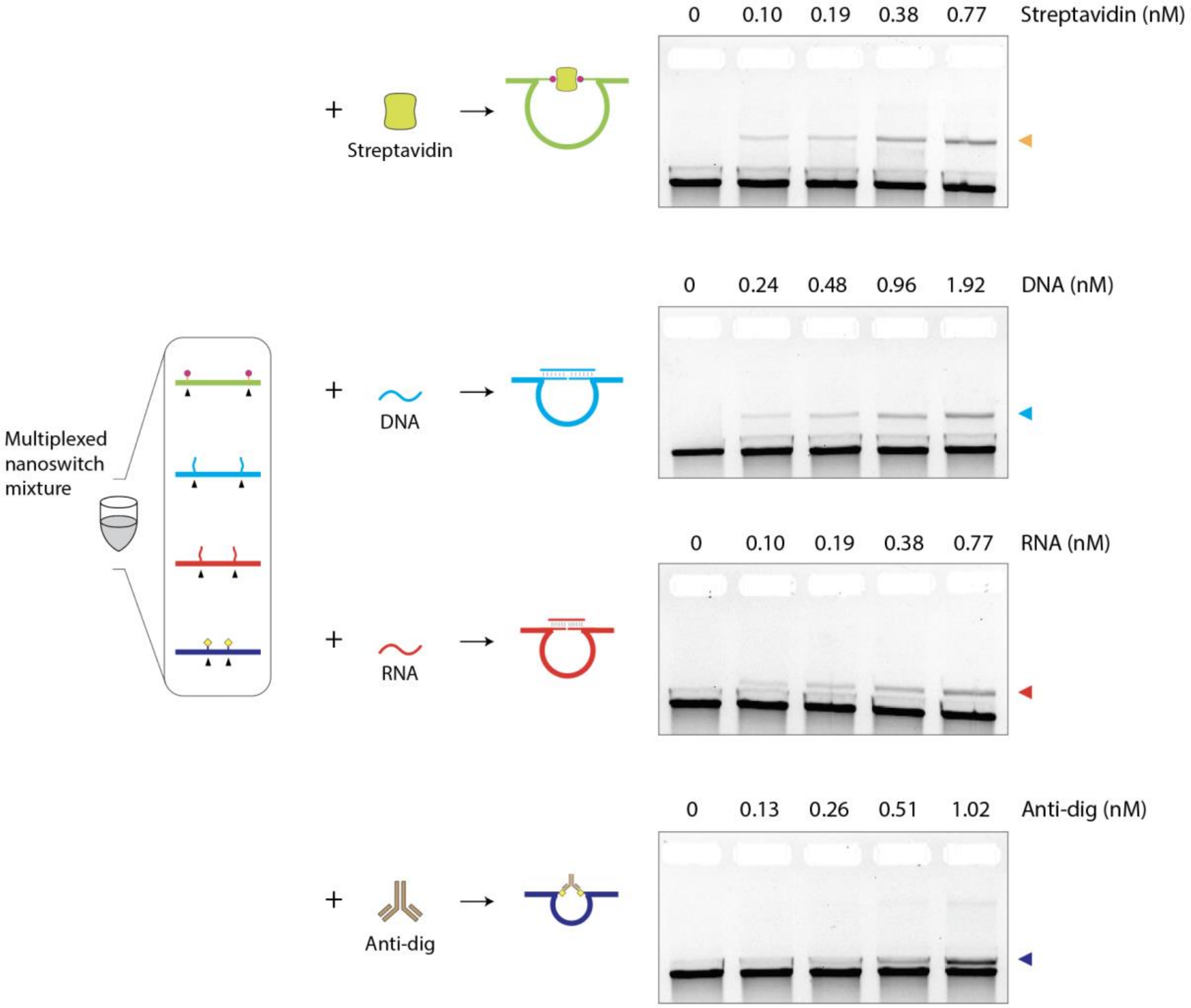


**Figure S9**. Concentration series of individual biomarkers using a nanoswitch mix.

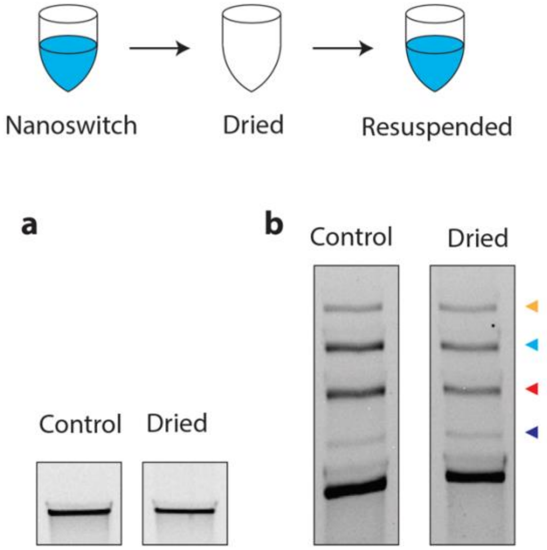


**Figure S10**. Stability and functionality of dried nanoswitches. (a) Nanoswitch mix is stable after drying and resuspending. (b) A dried and resuspended nanoswitch mixture can detect the four different target biomolecules.

Complete list of all sequences used. All sequences are written from 5' to 3'.

| Backbone oligonucleotides | | |
|---|---|---|
| **BB#** | **Sequence** | **Length** |
| 1 | AGAGCATAAAGCTAAATCGGTTGTACCAAAAACATTATGACCCTGTAATACTTTTGCGGG | 60 |
| 2 | AGAAGCCTTTATTTCAACGCAAGGATAAAAATTTTTAGAACCCTCATATATTTTAAATGC | 60 |
| 3 | AATGCCTGAGTAATGTGTAGGTAAAGATTCAAAAGGGTGAGAAAGGCCGGAGACAGTCAA | 60 |
| 4 | ATCACCATCAATATGATATTCAACCGTTCTAGCTGATAAATTAATGCCGGAGAGGGTAGC | 60 |
| 5 | TATTTTTGAGAGATCTACAAAGGCTATCAGGTCATTGCCTGAGAGTCTGGAGCAAACAAG | 60 |
| 6 | AGAATCGATGAACGGTAATCGTAAAACTAGCATGTCAATCATATGTACCCCGGTTGATAA | 60 |
| 7 | TCAGAAAAGCCCCAAAAACAGGAAGATTGTATAAGCAAATATTTAAATTGTAAACGTTAA | 60 |
| 8 | TATTTTGTTAAAATTCGCATTAAATTTTTGTTAAATCAGCTCATTTTTTAACCAATAGGA | 60 |
| 9 | ACGCCATCAAAAATAATTCGCGTCTGGCCTTCCTGTAGCCAGCTTTCATCAACATTAAAT | 60 |
| 10 | GGATAGGTCACGTTGGTGTAGATGGGCGCATCGTAACCGTGCATCTGCCAGTTTGAGGGG | 60 |
| 11 | ACGACGACAGTATCGGCCTCAGGAAGATCGCACTCCAGCCAGCTTTCCGGCACCGCTTCT | 60 |
| 12 | GGTGCCGGAAACCAGGCAAAGCGCCATTCGCCATTCAGGCTGCGCAACTGTTGGGAAGGG | 60 |
| 13 | CGATCGGTGCGGGCCTCTTCGCTATTACGCCAGCTGGCGAAAGGGGGATGTGCTGCAAGG | 60 |
| 14 | CGATTAAGTTGGGTAACGCCAGGGTTTTCCCAGTCACGACGTTGTAAAACGACGGCCAGT | 60 |
| 15 | GCCAAGCTTGCATGCCTGCAGGTCGACTCTAGAGGATCCCCGGGTACCGAGCTCGAATTC | 60 |
| 16 | GTAATCATGGTCATAGCTGTTTCCTGTGTGAAATTGTTATCCGCTCACAATTCCACACAA | 60 |
| 17 | CATACGAGCCGGAAGCATAAAGTGTAAAGCCTGGGGTGCCTAATGAGTGAGCTAACTCAC | 60 |
| 18 | ATTAATTGCGTTGCGCTCACTGCCCGCTTTCCAGTCGGGAAACCTGTCGTGCCAGCTGCA | 60 |
| 19 | TTAATGAATCGGCCAACGCGCGGGGAGAGGCGGTTTGCGTATTGGGCGCCAGGGTGGTTT | 60 |
| 20 | GTTGCAGCAAGCGGTCCACGCTGGTTTGCCCCAGCAGGCGAAAATCCTGTTTGATGGTGG | 60 |
| 21 | TTCCGAAATCGGCAAAATCCCTTATAAATCAAAAGAATAGCCCGAGATAGGGTTGAGTGT | 60 |
| 22 | TGTTCCAGTTTGGAACAAGAGTCCACTATTAAAGAACGTGGACTCCAACGTCAAAGGGCG | 60 |
| 23 | AAAAACCGTCTATCAGGGCGATGGCCCACTACGTGAACCATCACCCAAATCAAGTTTTTT | 60 |
| 24 | GGGGTCGAGGTGCCGTAAAGCACTAAATCGGAACCCTAAAGGGAGCCCCCGATTTAGAGC | 60 |
| 25 | TTGACGGGGAAAGCCGGCGAACGTGGCGAGAAAGGAAGGGAAGAAAGCGAAAGGAGCGGG | 60 |
| 26 | CGCTAGGGCGCTGGCAAGTGTAGCGGTCACGCTGCGCGTAACCACCACACCCGCCGCGCT | 60 |
| 27 | TAATGCGCCGCTACAGGGCGCGTACTATGGTTGCTTTGACGAGCACGTATAACGTGCTTT | 60 |
| 28 | CCTCGTTAGAATCAGAGCGGGAGCTAAACAGGAGGCCGATTAAAGGGATTTTAGACAGGA | 60 |
| 29 | ACGGTACGCCAGAATCCTGAGAAGTGTTTTTATAATCAGTGAGGCCACCGAGTAAAAGAG | 60 |
| 30 | TTGCCTGAGTAGAAGAACTCAAACTATCGGCCTTGCTGGTAATATCCAGAACAATATTAC | 60 |
| 31 | CGCCAGCCATTGCAACAGGAAAAACGCTCATGGAAATACCTACATTTTGACGCTCAATCG | 60 |
| 32 | TCTGAAATGGATTATTTACATTGGCAGATTCACCAGTCACACGACCAGTAATAAAAGGGA | 60 |
| 33 | CATTCTGGCCAACAGAGATAGAACCCTTCTGACCTGAAAGCGTAAGAATACGTGGCACAG | 60 |
| 34 | ACAATATTTTTGAATGGCTATTAGTCTTTAATGCGCGAACTGATAGCCCTAAAACATCGC | 60 |
| 35 | CATTAAAAATACCGAACGAACCACCAGCAGAAGATAAAACAGAGGTGAGGCGGTCAGTAT | 60 |
| 36 | TAACACCGCCTGCAACAGTGCCACGCTGAGAGCCAGCAGCAAATGAAAAATCTAAAGCAT | 60 |

| | | |
|---|---|---|
| 37 | CACCTTGCTGAACCTCAAATATCAAACCCTCAATCAATATCTGGTCAGTTGGCAAATCAA | 60 |
| 38 | CAGTTGAAAGGAATTGAGGAAGGTTATCTAAAATATCTTTAGGAGCACTAACAACTAATA | 60 |
| 39 | GATTAGAGCCGTCAATAGATAATACATTTGAGGATTTAGAAGTATTAGACTTTACAAACA | 60 |
| 40 | CATTATCATTTTGCGGAACAAAGAAACCACCAGAAGGAGCGGAATTATCATCATATTCCT | 60 |
| 41 | GATTATCAGATGATGGCAATTCATCAATATAATCCTGATTGTTTGGATTATACTTCTGAA | 60 |
| 42 | TAATGGAAGGGTTAGAACCTACCATATCAAAATTATTTGCACGTAAAACAGAAATAAAGA | 60 |
| 43 | AATTGCGTAGATTTTCAGGTTTAACGTCAGATGAATATACAGTAACAGTACCTTTTACAT | 60 |
| 44 | CGGGAGAAACAATAACGGATTCGCCTGATTGCTTTGAATACCAAGTTACAAAATCGCGCA | 60 |
| 45 | GAGGCGAATTATTCATTTCAATTACCTGAGCAAAAGAAGATGATGAAACAAACATCAAGA | 60 |
| 46 | AAACAAAATTAATTACATTTAACAATTTCATTTGAATTACCTTTTTTAATGGAAACAGTA | 60 |
| 47 | CATAAATCAATATATGTGAGTGAATAACCTTGCTTCTGTAAATCGTCGCTATTAATTAAT | 60 |
| 48 | TTTCCCTTAGAATCCTTGAAAACATAGCGATAGCTTAGATTAAGACGCTGAGAAGAGTCA | 60 |
| 49 | ATAGTGAATTTATCAAAATCATAGGTCTGAGAGACTACCTTTTTAACCTCCGGCTTAGGT | 60 |
| 50 | GAAAACTTTTTCAAATATATTTTAGTTAATTTCATCTTCTGACCTAAATTTAATGGTTTG | 60 |
| 51 | AAATACCGACCGTGTGATAAATAAGGCGTTAAATAAGAATAAACACCGGAATCATAATTA | 60 |
| 52 | CTAGAAAAAGCCTGTTTAGTATCATATGCGTTATACAAATTCTTACCAGTATAAAGCCAA | 60 |
| 53 | CGCTCAACAGTAGGGCTTAATTGAGAATCGCCATATTTAACAACGCCAACATGTAATTTA | 60 |
| 54 | GGCAGAGGCATTTTCGAGCCAGTAATAAGAGAATATAAAGTACCGACAAAAGGTAAAGTA | 60 |
| 55 | ATTCTGTCCAGACGACGACAATAAACAACATGTTCAGCTAATGCAGAACGCGCCTGTTTA | 60 |
| 56 | TCAACAATAGATAAGTCCTGAACAAGAAAAATAATATCCCATCCTAATTTACGAGCATGT | 60 |
| 57 | AGAAACCAATCAATAATCGGCTGTCTTTCCTTATCATTCCAAGAACGGGTATTAAACCAA | 60 |
| 58 | GTACCGCACTCATCGAGAACAAGCAAGCCGTTTTTATTTTCATCGTAGGAATCATTACCG | 60 |
| 59 | CGCCCAATAGCAAGCAAATCAGATATAGAAGGCTTATCCGGTATTCTAAGAACGCGAGGC | 60 |
| 60 | ATTTTGCACCCAGCTACAATTTTATCCTGAATCTTACCAACGCTAACGAGCGTCTTTCCA | 60 |
| 61 | GAGCCTAATTTGCCAGTTACAAAATAAACAGCCATATTATTTATCCCAATCCAAATAAGA | 60 |
| 62 | AACGATTTTTTGTTTAACGTCAAAAATGAAAATAGCAGCCTTTACAGAGAGAATAACATA | 60 |
| 63 | AAAACAGGGAAGCGCATTAGACGGGAGAATTAACTGAACACCCTGAACAAAGTCAGAGGG | 60 |
| 64 | TAATTGAGCGCTAATATCAGAGAGATAACCCACAAGAATTGAGTTAAGCCCAATAATAAG | 60 |
| 65 | AGCAAGAAACAATGAAATAGCAATAGCTATCTTACCGAAGCCCTTTTTAAGAAAAGTAAG | 60 |
| 66 | CAGATAGCCGAACAAAGTTACCAGAAGGAAACCGAGGAAACGCAATAATAACGGAATACC | 60 |
| 67 | CAAAAGAACTGGCATGATTAAGACTCCTTATTACGCAGTATGTTAGCAAACGTAGAAAAT | 60 |
| 68 | ACATACATAAAGGTGGCAACATATAAAAGAAACGCAAAGACACCACGGAATAAGTTTATT | 60 |
| 69 | TTGTCACAATCAATAGAAAATTCATATGGTTTACCAGCGCCAAAGACAAAAGGGCGACAT | 60 |
| 70 | TCACCGTCACCGACTTGAGCCATTTGGGAATTAGAGCCAGCAAAATCACCAGTAGCACCA | 60 |
| 71 | TTACCATTAGCAAGGCCGGAAACGTCACCAATGAAACCATCGATAGCAGCACCGTAATCA | 60 |
| 72 | GTAGCGACAGAATCAAGTTTGCCTTTAGCGTCAGACTGTAGCGCGTTTTCATCGGCATTT | 60 |
| 73 | TCGGTCATAGCCCCCTTATTAGCGTTTGCCATCTTTTCATAATCAAAATCACCGGAACCA | 60 |
| 74 | GAGCCACCACCGGAACCGCCTCCCTCAGAGCCGCCACCCTCAGAACCGCCACCCTCAGAG | 60 |
| 75 | CCACCACCCTCAGAGCCGCCACCAGAACCACCACCAGAGCCGCCGCCAGCATTGACAGGA | 60 |
| 76 | GGTTGAGGCAGGTCAGACGATTGGCCTTGATATTCACAAACAAATAAATCCTCATTAAAG | 60 |
| 77 | CCAGAATGGAAAGCGCAGTCTCTGAATTTACCGTTCCAGTAAGCGTCATACATGGCTTTT | 60 |

| | | |
|---|---|---|
| 78 | GATGATACAGGAGTGTACTGGTAATAAGTTTTAACGGGGTCAGTGCCTTGAGTAACAGTG | 60 |
| 79 | CCCGTATAAACAGTTAATGCCCCCTGCCTATTTCGGAACCTATTATTCTGAAACATGAAA | 60 |
| 80 | CCAGGCGGATAAGTGCCGTCGAGAGGGTTGATATAAGTATAGCCCGGAATAGGTGTATCA | 60 |
| 81 | CCGTACTCAGGAGGTTTAGTACCGCCACCCTCAGAACCGCCACCCTCAGAACCGCCACCC | 60 |
| 82 | TCAGAGCCACCACCCTCATTTTCAGGGATAGCAAGCCCAATAGGAACCCATGTACCGTAA | 60 |
| 83 | CACTGAGTTTCGTCACCAGTACAAACTACAACGCCTGTAGCATTCCACAGACAGCCCTCA | 60 |
| 84 | TAGTTAGCGTAACGATCTAAAGTTTTGTCGTCTTTCCAGACGTTAGTAAATGAATTTTCT | 60 |
| 85 | GTATGGGATTTTGCTAAACAACTTTCAACAGTTTCAGCGGAGTGAGAATAGAAAGGAACA | 60 |
| 86 | ACTAAAGGAATTGCGAATAATAATTTTTTCACGTTGAAAATCTCCAAAAAAAAGGCTCCA | 60 |
| 87 | AAAGGAGCCTTTAATTGTATCGGTTTATCAGCTTGCTTTCGAGGTGAATTTCTTAAACAG | 60 |
| 88 | CTTGATACCGATAGTTGCGCCGACAATGACAACAACCATCGCCCACGCATAACCGATATA | 60 |
| 89 | TTCGGTCGCTGAGGCTTGCAGGGAGTTAAAGGCCGCTTTTGCGGGATCGTCACCCTCAGC | 60 |
| 90 | CTTTTTCATGAGGAAGTTTCCATTAAACGGGTAAAATACGTAATGCCACTACGAAGGCAC | 60 |
| 91 | CAACCTAAAACGAAAGAGGCAAAAGAATACACTAAAACACTCATCTTTGACCCCCAGCGA | 60 |
| 92 | TTATACCAAGCGCGAAACAAAGTACAACGGAGATTTGTATCATCGCCTGATAAATTGTGT | 60 |
| 93 | CGAAATCCGCGACCTGCTCCATGTTACTTAGCCGGAACGAGGCGCAGACGGTCAATCATA | 60 |
| 94 | AGGGAACCGAACTGACCAACTTTGAAAGAGGACAGATGAACGGTGTACAGACCAGGCGCA | 60 |
| 95 | TAGGCTGGCTGACCTTCATCAAGAGTAATCTTGACAAGAACCGGATATTCATTACCCAAA | 60 |
| 96 | TCAACGTAACAAAGCTGCTCATTCAGTGAATAAGGCTTGCCCTGACGAGAAACACCAGAA | 60 |
| 97 | CGAGTAGTAAATTGGGCTTGAGATGGTTTAATTTCAACTTTAATCATTGTGAATTACCTT | 60 |
| 98 | ATGCGATTTTAAGAACTGGCTCATTATACCAGTCAGGACGTTGGGAAGAAAAATCTACGT | 60 |
| 99 | TAATAAAACGAACTAACGGAACAACATTATTACAGGTAGAAAGATTCATCAGTTGAGATT | 60 |
| 100 | TAAGAGCAACACTATCATAACCCTCGTTTACCAGACGACGATAAAAACCAAAATAGCGAG | 60 |
| 101 | AGGCTTTTGCAAAAGAAGTTTTGCCAGAGGGGGTAATAGTAAAATGTTTAGACTGGATAG | 60 |
| 102 | CGTCCAATACTGCGGAATCGTCATAAATATTCATTGAATCCCCCTCAAATGCTTTAAACA | 60 |
| 103 | GTTCAGAAAACGAGAATGACCATAAATCAAAAATCAGGTCTTTACCCTGACTATTATAGT | 60 |
| 104 | CAGAAGCAAAGCGGATTGCATCAAAAAGATTAAGAGGAAGCCCGAAAGACTTCAAATATC | 60 |
| 105 | GCGTTTTAATTCGAGCTTCAAAGCGAACCAGACCGGAAGCAAACTCCAACAGGTCAGGAT | 60 |
| 106 | TAGAGAGTACCTTTAATTGCTCCTTTTGATAAGAGGTCATTTTTGCGGATGGCTTAGAGC | 60 |
| 107 | TTAATTGCTGAATATAATGCTGTAGCTCAACATGTTTTAAATATGCAACTAAAGTACGGT | 60 |
| 108 | GTCTGGAAGTTTCATTCCATATAACAGTTGATTCCCAATTCTGCGAACGAGTAGATTTAG | 60 |
| 109 | TTTGACCATTAGATACATTTCGCAAATGGTCAATAACCTGTTTAGCTAT | 49 |

Regions in **bold** indicate locations where detector strands are placed. Blue and green regions are single stranded extensions on detectors that are complementary to two halves of the input strands (color coded in target sequences). For nanoswitch construction and detector placement, refer to Figure S1.

| Variable sequences | | |
|---|---|---|
| # | Sequence | Length |
| V1 | AACATCCAATAAATCATACAGGCAAGGCAAAGAATTAGCAAAATTAAGCAATAAAGCCTC | 60 |
| V2 | GTGAGCGAGTAACAACCCGTCGGATTCTCCGTGGGAACAAACGGCGGATTGACCGTAATG | 60 |
| V3 | TTCTTTTCACCAGTGAGACG**GGCAACAGCTGATTGCCCTTCACCGCCTGGCCCTGAGAGA** | 60 |
| V4 | TCTGTCCATCACGCAAATTA**ACCGTTGTAGCAATACTTCTTTGATTAGTAATAACATCAC** | 60 |
| V5 | **ATTCGACAACTCGTATTAAATCCTTTGCCCGAACGTTATT**AATTTTAAAAGTTTGAGTAA | 60 |
| V6 | **TGGGTTATATAACTATATGTAAATGCTGATGCAAATCCAA**TCGCAAGACAAAGAACGCGA | 60 |
| V7 | **GTTTTAGCGAACCTCCCGACTTGCGGGAGGTTTTGAAGCC**TTAAATCAAGATTAGTTGCT | 60 |
| V8 | **TCAACCGATTGAGGGAGGGAAGGTAAATATTGACGGAAAT**TATTCATTAAAGGTGAATTA | 60 |
| V9 | **GTATTAAGAGGCTGAGACTCCTCAAGAGAAGGATTAGGAT**TAGCGGGGTTTTGCTCAGTA | 60 |
| V10 | **AGCGAAAGACAGCATCGGAACGAGGGTAGCAACGGCTACA**GAGGCTTTGAGGACTAAAGA | 60 |
| V11 | TAGGAATACCACATTCAACTAATGCAGATACATAACGCCAAAAGGAATTACGAGGCATAG | 60 |
| V12 | ATTTTCATTTGGGGCGCGAGCTGAAAAGGTGGCATCAATTCTACTAATAGTAGTAGCATT | 60 |

| Detector sequences for characterization (4-8 loop size) | | |
|---|---|---|
| # | Sequence | Length |
| V4 SP 40-15 | ACCGTTGTAGCAATACTTCTTTGATTAGTAATAACATCACTCACAATTAGATCCT | 55 |
| V8 SP 15-40 | GTAACTTACACATGATCAACCGATTGAGGGAGGGAAGGTAAATATTGACGGAAAT | 55 |
| V4 CF 40-12 | ACCGTTGTAGCAATACTTCTTTGATTAGTAATAACATCACGTCGTCTGCTGC | 52 |
| V8 CF 12-40 | TGCTGCTGCTGCTCAACCGATTGAGGGAGGGAAGGTAAATATTGACGGAAAT | 52 |
| V8 CF 11-40 | TGCTGCTGCTGTCAACCGATTGAGGGAGGGAAGGTAAATATTGACGGAAAT | 51 |
| V8 CF 10-40 | TGCTGCTGCTTCAACCGATTGAGGGAGGGAAGGTAAATATTGACGGAAAT | 50 |
| V8 CF 9-40 | TGCTGCTGCTCAACCGATTGAGGGAGGGAAGGTAAATATTGACGGAAAT | 49 |
| V4 TS 40-11 | ACCGTTGTAGCAATACTTCTTTGATTAGTAATAACATCACGAACCGTATAT | 51 |
| V8 TS 10-40 | CCTATGGCCCTCAACCGATTGAGGGAGGGAAGGTAAATATTGACGGAAAT | 50 |
| V4 BRCA 40-13 | ACCGTTGTAGCAATACTTCTTTGATTAGTAATAACATCACCTTCCAACAGCTA | 53 |
| V8 BRCA 12-40 | TAAACAGTCCTGTCAACCGATTGAGGGAGGGAAGGTAAATATTGACGGAAAT | 52 |
| V4 HIV 40-12 | ACCGTTGTAGCAATACTTCTTTGATTAGTAATAACATCACAGTCAGTGTGGA | 52 |
| V8 HIV 11-40 | AAATCTCTAGCTCAACCGATTGAGGGAGGGAAGGTAAATATTGACGGAAAT | 51 |
| V4 WS 40-17 | ACCGTTGTAGCAATACTTCTTTGATTAGTAATAACATCACCATCTTCAAATCCATCT | 57 |
| V8 WS 17-40 | TCTTTTCATTCCACTTTTCAACCGATTGAGGGAGGGAAGGTAAATATTGACGGAAAT | 57 |

| Detector sequences for gene barcodes (different loop sizes) | | |
|---|---|---|
| Name | Sequence | Length |
| V4 SP 40-15 | ACCGTTGTAGCAATACTTCTTTGATTAGTAATAACATCACTCACAATTAGATCCT | 55 |
| V5 SP 15-40 | GTAACTTACACATGAATTCGACAACTCGTATTAAATCCTTTGCCCGAACGTTATT | 55 |
| V4 CF 40-12 | ACCGTTGTAGCAATACTTCTTTGATTAGTAATAACATCACGTCGTCTGCTGC | 52 |
| V6 CF 12-40 | TGCTGCTGCTGCTGGGTTATATAACTATATGTAAATGCTGATGCAAATCCAA | 52 |
| V4 TS 40-11 | ACCGTTGTAGCAATACTTCTTTGATTAGTAATAACATCACGAACCGTATAT | 51 |
| V7 TS 10-40 | CCTATGGCCCGTTTTAGCGAACCTCCCGACTTGCGGGAGGTTTTGAAGCC | 50 |
| V4 BRCA 40-13 | ACCGTTGTAGCAATACTTCTTTGATTAGTAATAACATCACCTTCCAACAGCTA | 53 |
| V8 BRCA 12-40 | TAAACAGTCCTGTCAACCGATTGAGGGAGGGAAGGTAAATATTGACGGAAAT | 52 |
| V4 HIV 40-12 | ACCGTTGTAGCAATACTTCTTTGATTAGTAATAACATCACAGTCAGTGTGGA | 52 |
| V9 HIV 11-40 | AAATCTCTAGCGTATTAAGAGGCTGAGACTCCTCAAGAGAAGGATTAGGAT | 51 |
| V3 WS 40-17 | GGCAACAGCTGATTGCCCTTCACCGCCTGGCCCTGAGAGACATCTTCAAATCCATCT | 57 |
| V10 WS 17-40 | TCTTTTCATTCCACTTTAGCGAAAGACAGCATCGGAACGAGGGTAGCAACGGCTACA | 57 |

| Detector sequences for mixed multiplexing barcodes (different loop sizes) | | |
|---|---|---|
| Name | Sequence | Length |
| V4 dig | ACCGTTGTAGCAATACTTCTTTGATTAGTAATAACATCAC-dig | 40 |
| V5 dig | dig-ATTCGACAACTCGTATTAAATCCTTTGCCCGAACGTTATT | 40 |
| V4 RNA | ACCGTTGTAGCAATACTTCTTTGATTAGTAATAACATCACCGCCAATATTT | 51 |
| V6 RNA | ACGTGCTGCTATGGGTTATATAACTATATGTAAATGCTGATGCAAATCCAA | 52 |
| V4 DNA | ACCGTTGTAGCAATACTTCTTTGATTAGTAATAACATCACCCAACAACAT | 50 |
| V7 DNA | GAAACTACCTAGTTTTAGCGAACCTCCCGACTTGCGGGAGGTTTTGAAGCC | 51 |
| V4 biotin | ACCGTTGTAGCAATACTTCTTTGATTAGTAATAACATCAC-biotin | 40 |
| V8 biotin | biotin-TCAACCGATTGAGGGAGGGAAGGTAAATATTGACGGAAAT | 40 |
| V4 miR-30c | ACCGTTGTAGCAATACTTCTTTGATTAGTAATAACATCACTCCAACACTGT | 51 |
| V5 miR-30c | ACTGGAAGATGATTCGACAACTCGTATTAAATCCTTTGCCCGAACGTTATT | 51 |
| V4 miR-141 | ACCGTTGTAGCAATACTTCTTTGATTAGTAATAACATCACGCTGAGAGTGTA | 52 |
| V7 miR-141 | GGATGTTTACAGTTTTAGCGAACCTCCCGACTTGCGGGAGGTTTTGAAGCC | 51 |
| BB37 PSA1 | CAATCAATATCTGGTCAGTTGGCAAATCAA-ab1 | 30 |
| V7 PSA2 | ab2-GTTTTAGCGAACCTCCCGACTTGCGGGAGG | 30 |

| Filler sequences | | |
|---|---|---|
| **Name** | **Sequence** | **Length** |
| V3 filler | TTCTTTTCACCAGTGAGACG | 20 |
| V4 filler | TCTGTCCATCACGCAAATTA | 20 |
| V5 filler | AATTTTAAAAGTTTGAGTAA | 20 |
| V6 filler | TCGCAAGACAAAGAACGCGA | 20 |
| V7 filler | TTAAATCAAGATTAGTTGCT | 20 |
| V8 filler | TATTCATTAAAGGTGAATTA | 20 |
| V9 filler | TAGCGGGGTTTTGCTCAGTA | 20 |
| V10 filler | GAGGCTTTGAGGACTAAAGA | 20 |
| BB37 filler (PSA) | CACCTTGCTGAACCTCAAATATCAAACCCT | 30 |
| V7 filler (PSA) | TTTTGAAGCCTTAAATCAAGATTAGTTGCT | 30 |

| Target sequences | | |
|---|---|---|
| **Name** | **Sequence** | **Length** |
| SP DNA | TCATGTGTAAGTTACAGGATCTAATTGTGA | 30 |
| CF DNA | GCAGCAGCAGCAGCAGCAGACGAC | 24 |
| TS DNA | GGGCCATAGGATATACGGTTC | 21 |
| BRCA1 DNA | CAGGACTGTTTATAGCTGTTGGAAG | 25 |
| HIV1 DNA | GCTAGAGATTTTCCACACTGACT | 23 |
| WS DNA | AAAGTGGAATGAAAAGAAGATGGATTTGAAGATG | 34 |
| CF 1 mismatch | GCAGCACCAGCAGCAGCAGACGAC | 24 |
| CF 2 mismatch | GCTGCACCAGCAGCAGCAGACGAC | 24 |
| CF 3 mismatch | GCTGGACCAGCAGCAGCAGACGAC | 24 |
| RNA (Fig 4) | UAGCAGCACGUAAAUAUUGGCG | 22 |
| DNA (Fig 4) | TAGGTAGTTTCATGTTGTTGG | 21 |
| miR-141 DNA | CATCTTCCAGTACAGTGTTGGA | 22 |
| miR-30c DNA | TGTAAACATCCTACACTCTCAGC | 23 |

| Other strands | | |
|---|---|---|
| **Name** | **Sequence** | **Length** |
| Blocking oligos | ACGGTCTCATGGCCCTTCAATC | 22 |
| BtsCI cut site oligo | CTACTAATAGTAGTAGCATTAACATCCAATAAATCATACA | 40 |
